# Functional connectivity correlates of the hierarchical p-factor model in youth at neurodevelopmental risk

**DOI:** 10.64898/2026.05.03.722505

**Authors:** Rebeca Ianov Vitanov, Danyal Akarca, CALM Team, Sarah E. Morgan, Jonathan S. Jones

**Author notes:** Correspondence: Rebeca Ianov Vitanov, MRC Cognition and Brain Sciences Unit University of Cambridge, Cambridge, United Kingdom. School of Biomedical Engineering and Imaging Sciences, King’s College London, London, United Kingdom (Sarah E. Morgan). Department of Electrical and Electronic Engineering, Imperial College London, London, United Kingdom (Danyal Akarca). These authors contributed equally. These authors jointly supervised this work (Sarah E. Morgan, Jonathan S. Jones).

## Abstract

**Background:** Emotional and cognitive difficulties often co-occur in neurodevelopmental conditions. While transdiagnostic, dimensional approaches offer a more precise framework for understanding mental health than diagnostic categories, their neural correlates in youth with learning difficulties remain poorly understood. This study investigates associations between transdiagnostic mental health dimensions and resting-state functional connectivity in struggling learners.

**Methods:** Cross-sectional behavioural data from the Centre for Attention, Learning and Memory (CALM) for struggling learners (N = 378) was used to replicate a hierarchical model of mental health from the Conners’ Parent Rating Short Form, the Revised Children’s Anxiety and Depression Scale and the Strengths and Difficulties Questionnaire. Functional connectomes were derived from resting-state fMRI data (N = 67), and partial least squares regression related mental health dimensions to connectivity within and between large-scale brain networks.

**Results:** The replicated model comprised a general p-factor, two broad domains (internalising and externalising), and three specific dimensions (specific internalising, neurodevelopmental and social maladjustment). Symptom severity was associated with two connectivity patterns: greater default mode network coupling to frontoparietal and attention networks, and reduced connectivity between visual and somatomotor systems. These effects were strongest for the neurodevelopmental and social maladjustment dimensions, respectively.

**Conclusions:** These findings align with population-level evidence linking mental health dimensions to brain network organization, extending it to struggling learners and offering new insight into the neural basis of mental health vulnerability in neurodevelopmentally at-risk youth.

## Introduction

Emotional and cognitive difficulties frequently co-occur during childhood and adolescence, particularly among youth with neurodevelopmental challenges (Murphy et al., 2015). These overlapping symptoms are associated with cognitive, social, sensory and motor difficulties (Morris-Rosendahl and Crocq, 2022; Hansen et al., 2018; Munir, 2016) and highlight limitations of traditional diagnostic categories. As a result, transdiagnostic approaches that capture shared dimensions of mental health have gained increasing attention. Mental health difficulties in childhood can affect emotional regulation and school performance, with early symptoms strongly predicting later executive functioning and academic outcomes (Donati et al., 2021; Agnafors et al., 2021; Milienos et al., 2021).

Given the limitations of traditional diagnostic categories based on DSM-V (APA, 2022) in capturing the heterogenous and overlapping symptoms of affective and neurodevelopmental conditions (Kapadia, Desai & Parikh, 2020; Insel, Cuthbert & Garvey, 2010; Wakefield, 2015), transdiagnostic approaches have emerged. These approaches focus on shared mechanisms and symptoms across disorders, often reflecting extensive comorbidity. Frameworks such as the Research Domain Criteria (RDoC) and Hierarchical Taxonomy of Psychopathology (HiTOP) highlight the common biological pathways and cognitive dysfunctions underlying many conditions (Caspi et al., 2014; Kotov et al., 2017; Lahey et al., 2021; Cuthbert, 2022). RDoC emphasizes six major neurobehavioral domains (Cuthbert, 2014), while HiTOP organizes symptoms hierarchically, from broad to specific dimensions (Kotov et al., 2022). For example, Holmes et al. (2021) identified hierarchically arranged mental health dimensions in struggling learners from the Centre for Attention, Learning and Memory (CALM) cohort. The model revealed a general p-factor at the apex and specific dimensions that predicted concurrent social, clinical, and educational outcomes, with the neurodevelopmental dimension being the strongest predictor of learning outcomes (Holmes et al., 2021).

Childhood and adolescence bring significant cognitive and emotional changes, underpinned by both structural and functional brain developments (Bethlehem et al., 2022; Grydeland et al., 2019; Supekar et al., 2009). Large-scale functional brain networks (also known as intrinsic connectivity networks, ICNs) undergo a developmental shift, with weakening short-range and strengthening long-range functional connectivity (FC), which are linked to later cognitive and mental health outcomes (Cao et al., 2014; Jones et al., 2022).

Deviations from typical connectivity, such as reduced within-network default-mode and increased default-mode to task-positive network connectivity, have been associated with cognitive impairments in at-risk youth (Nomi and Uddin, 2015; Sripada et al., 2014).

Studies have also connected hierarchical models of mental health to patterns of ICN connectivity, offering insights for clinical interventions (Xia et al., 2018; Karcher et al., 2021). For instance, Karcher et al. (2021) found associations between the general psychopathology (‘p’) dimensions, neurodevelopmental symptoms, and altered default-mode network (DMN) functional connectivity in the Adolescent Brain Cognitive Development (ABCD) cohort (Casey et al., 2018). Similarly, research from the Philadelphia Neurodevelopmental Cohort (PNC; Satterthwaite et al., 2014) linked mental health dimensions such as mood, psychosis, fear, and externalizing symptoms to increased integration between the DMN and executive networks, with internalizing symptoms tied to enhanced functional connectivity between ventral attention and salience networks (Xia et al., 2018). Mapping behavioural outcomes in childhood and adolescence to brain connectivity thus offers a valuable approach for identifying neurobiological mechanisms underlying cognitive and mental health differences. While the large, community-based cohorts such as the PNC and the ABCD study are invaluable for identifying population-level patterns of brain-behaviour relationships, they primarily include typically developing children and therefore underrepresent those with significant learning or attentional difficulties. As a result, they offer limited insight into the neurobiological mechanisms underpinning mental health problems in youth at neurodevelopmental risk.

While previous studies within the CALM cohort have examined associations between externalising behaviours and functional brain connectivity (Bathelt et al., 2018; Astle et al., 2019; Jones et al., 2021; Monaghan et al., 2026), the current study advances this work by investigating how a hierarchical, dimensional model of mental health – spanning both internalising and externalising symptoms – relates to FC in this transdiagnostic sample of struggling learners. The CALM cohort represents children at neurodevelopmental risk, referred for difficulties in attention, learning, or memory by health or educational professionals. This heterogeneous group, which includes individuals both below and above clinical thresholds, provides a unique opportunity to characterise the relationship between mental health dimensions and brain network organisation across a broad spectrum of cognitive difficulties, offering a transdiagnostic perspective on the neural basis of mental health in neurodivergent youth.

## Methods

### Behavioural data

#### Participant characteristics

The data used in this study was derived from the Centre for Attention, Learning and Memory (CALM cohort; Holmes et al., 2019). The CALM subset of participants at neurodevelopmental risk included 805 children for whom parent-reported behavioural data was available (CALM800: 69% male, mean age = 9.48 years; standard deviation of age = 2.38 years). Of these, 63% were referred by education practitioners (e.g., educational psychologist, special educational needs coordinator), 33% were referred by health professionals (e.g., clinical psychologist, child psychiatrist) and 4% by speech and language therapists. The participants at neurodevelopmental risk were referred as having at least one difficulty in attention, memory, language, literacy and mathematics or poor school progress. Forty percent of the cohort had an official diagnosis and 8% had at least two diagnoses. The most common diagnoses were attention deficit hyperactivity disorder (ADHD; N = 197), autism spectrum disorder (ASD; N = 57) and learning disorders (e.g. dyslexia, dyscalculia, developmental language disorder, N = 62).

#### Behavioural measures

The behavioural variables were derived from three widely used parent-report measures capturing internalising and externalising symptoms: the Revised Child Anxiety and Depression Scale-Parent Version (RCADS; Chorpita et al., 2000), Conners-3 Parent Short Form (Conners, 2008) and Strengths and Difficulties Questionnaire (SDQ; Goodman & Goodman, 2009). The included subscales were:

- RCADS: Separation anxiety, social phobia, generalised anxiety, panic disorder, obsessive–compulsive disorder, depression
- Conners: Inattention, hyperactivity/impulsivity, executive function, aggression, peer relations, learning problems
- SDQ: Prosocial behaviour (reverse-coded) and conduct problems

The choice to include these specific subscales was motivated by the aim to maximise the range of symptoms captured, while including only a single indicator of any particular symptom from all subscales. The current study included all Conners’ questionnaire subscales, including the ‘Learning problems’ subscale, which Holmes et al. (2021) excluded as it was used as a dependent variable when predicting concurrent learning outcomes. Age and sex effects were removed from the raw mental health subscale scores using linear regression to ensure that these potential confounders did not influence the results.

A total of N = 771 participants had SDQ and Conners data available, and N = 378 had additional RCADS data (the RCADS was introduced later in recruitment). Participants lacking data from one or more included subscales or older than 16 years were excluded. A six-dimensions hierarchical model was derived from participants with complete SDQ, Conners and RCADS data. To maximise the statistical power obtained from the larger number of subjects with SDQ and Conners data, but without RCADS data, these subjects (N = 771, the ‘SDQ and Conners cohort’) were used to derive the externalising mental health dimensions (externalising, neurodevelopmental and social maladjustment). The participants with SDQ, Conners and RCADS questionnaire data available (N = 378, the ‘RCADS cohort’) were used to derive the general p-factor, the internalising and specific internalising dimensions. The number of dimensions was determined by parallel analysis (Horn, 1965) and the hierarchical structure was extracted using Goldberg’s bass-ackward method (Goldberg, 2006) with geomin oblique rotations, following Holmes et al. (2021).

#### Participant subsets for brain–behaviour analyses

Of the N = 771 participants with SDQ and Conners data, N = 171 also had rs-fMRI data (‘SDQ & Conners FC subset’); of the N = 378 participants with all three questionnaires, N = 67 had rs-fMRI data (‘RCADS FC subset’). Participants older than 17 years at the time of the MRI scan (which took place a few months later than the behavioural testing) were excluded from the analysis. No significant differences in questionnaire subscales were observed between the two FC subsets (Table 1).

**Table 1.** Raw scores for the subscales included from each of the three mental health questionnaires (SDQ, Conners and RCADS). The mean value (M) and standard deviation (SD) are shown for scores in each of the two subsets used to relate mental health to rs-fMRI brain connectivity: the ‘SDQ & Conners FC subset’ (N = 171) and the ‘RCADS FC subset’ (N = 67). The differences between the two groups, for each questionnaire subscale, were not significant (Mann-Whitney test p-value > 0.05). The age range was 6.16 - 16.3 years for the ‘SDQ & Conners FC subset’, and 7.6 - 16.1 years old for the ‘RCADS FC subset’. The former subset (N = 171) consisted of 112 female and 59 male participants, whereas the latter subset (N = 67) comprised of 15 female and 52 male participants. SDQ = Strengths and Difficulties Questionnaire; Conners’ = Conners-3 Parent Rating Scale Short Form; RCADS = Revised Child and Anxiety and Depression Scale (Parent Version); GAD = Generalised anxiety disorder; OCD = Obsessive compulsive disorder.

| Subscale/Academic Test<br>raw scores | SDQ & Conners FC<br>subset<br>(N = 171) |  | RCADS FC subset<br>(N = 67) |  | P-values |
| --- | --- | --- | --- | --- | --- |
|  | M | SD | M | SD |  |
| SDQ Conduct | 3.08 | 2.24 | 3.29 | 2.21 | 0.6 |
| SDQ Prosocial | 7.05 | 2.36 | 6.55 | 2.29 | 0.24 |
| Conners' Inattention | 11.44 | 3.65 | 11.32 | 3.69 | 0.8 |
| Conners' Hyperactivity/<br>Impulsivity | 9.65 | 5.61 | 10.07 | 5.29 | 0.5 |
| Conners' Executive<br>Function | 10.20 | 3.71 | 10.52 | 3.84 | 0.5 |
| Conners' Aggression | 2.98 | 3.52 | 3.23 | 3.41 | 0.68 |
| Conners' Peer Relations | 5.48 | 4.71 | 6.14 | 4.57 | 0.31 |
| RCADS Separation anxiety | - | - | 5.07 | 3.87 | - |
| RCADS GAD | - | - | 5.98 | 4.18 | - |
| RCADS Panic | - | - | 3.56 | 3.47 | - |
| RCADS Social | - | - | 12.64 | 6.63 | - |
| RCADS OCD | - | - | 2.41 | 2.56 | - |
| RCADS Depression | - | - | 9.10 | 5.65 | - |

### Resting-state fMRI

#### fMRI data preprocessing

All scans were obtained on a Siemens 3T Prisma-fit system (Siemens Healthcare, Erlangen, Germany), using a 32-channel quadrature head coil at the MRC Cognition and Brain Sciences Unit, University of Cambridge. For the resting-state fMRI, 270 T2*-weighted whole-brain echo planar images (EPIs) were acquired over nine minutes (repetition time [TR] = 2s; echo time [TE] = 30ms; flip angle = 78 degrees, 3×3×3mm). The first 4 volumes were discarded to ensure steady state magnetization. Participants were instructed to lie still with their eyes closed and to not fall asleep. For registration of functional images, T1-weighted volume scans were acquired using a whole-brain coverage 3D Magnetization Prepared Rapid Acquisition Gradient Echo (MP-RAGE) sequence acquired using 1-mm isometric image resolution (TR = 2.25s, TE = 2.98ms, flip angle = 9 degrees, 1x1x1mm). Further details about the data acquisition and pre-processing can be found in the Supplementary Methods (Resting-state functional MRI data) and Jones et al., 2021. Briefly, the fMRI data were minimally pre-processed using fMRIPrep 1.5.0 for slice-timing correction, realignment, co-registration, segmentation, and normalization. Denoising was performed with fmridenoise, including band-pass filtering (0.01–0.1 Hz), 24 head motion parameters, 10 aCompCor components, motion spike regressors (> 0.5 mm displacement), and trend removal. Participants with excessive motion (mean framewise displacement > 0.5 mm or > 20% spikes) were excluded, yielding a final sample of N = 241 children with useable fMRI data.

#### Functional connectome construction and further preprocessing

The denoised fMRI data was parcellated according to a 274 region resting-state fMRI cortical and subcortical parcellation based on the Brainnetome atlas (Fan et al., 2016).

All cerebellar regions were excluded, resulting in 246 regions (210 cortical and 36 subcortical regions; see Figure 1a). Pearson correlations were computed pairwise between the regional time-series for each individual, generating an individual 246 x 246 connectivity matrix per subject, as shown in Figure 1b. Proportional thresholding was used to remove spurious false-positive edges. This method allows the retention of an arbitrary top % of the strongest connection weights, controlling for the number of connections across individuals, while removing noisy edges (van den Heuvel et al., 2008). The negative edges in the unthresholded weighted connectivity matrices were converted to 0 and proportional thresholding was then performed for a range of six thresholds: 5%, 10%, 15%, 20%, 25%, 30%. This resulted in connectivity matrices with e.g. 15068 edges at the 25% threshold. Results reported throughout the rest of the study were obtained for the 25% connectome threshold. We tested whether results were consistent across thresholds and the other thresholds also resulted in significant results unless otherwise stated.

**Figure 1:**
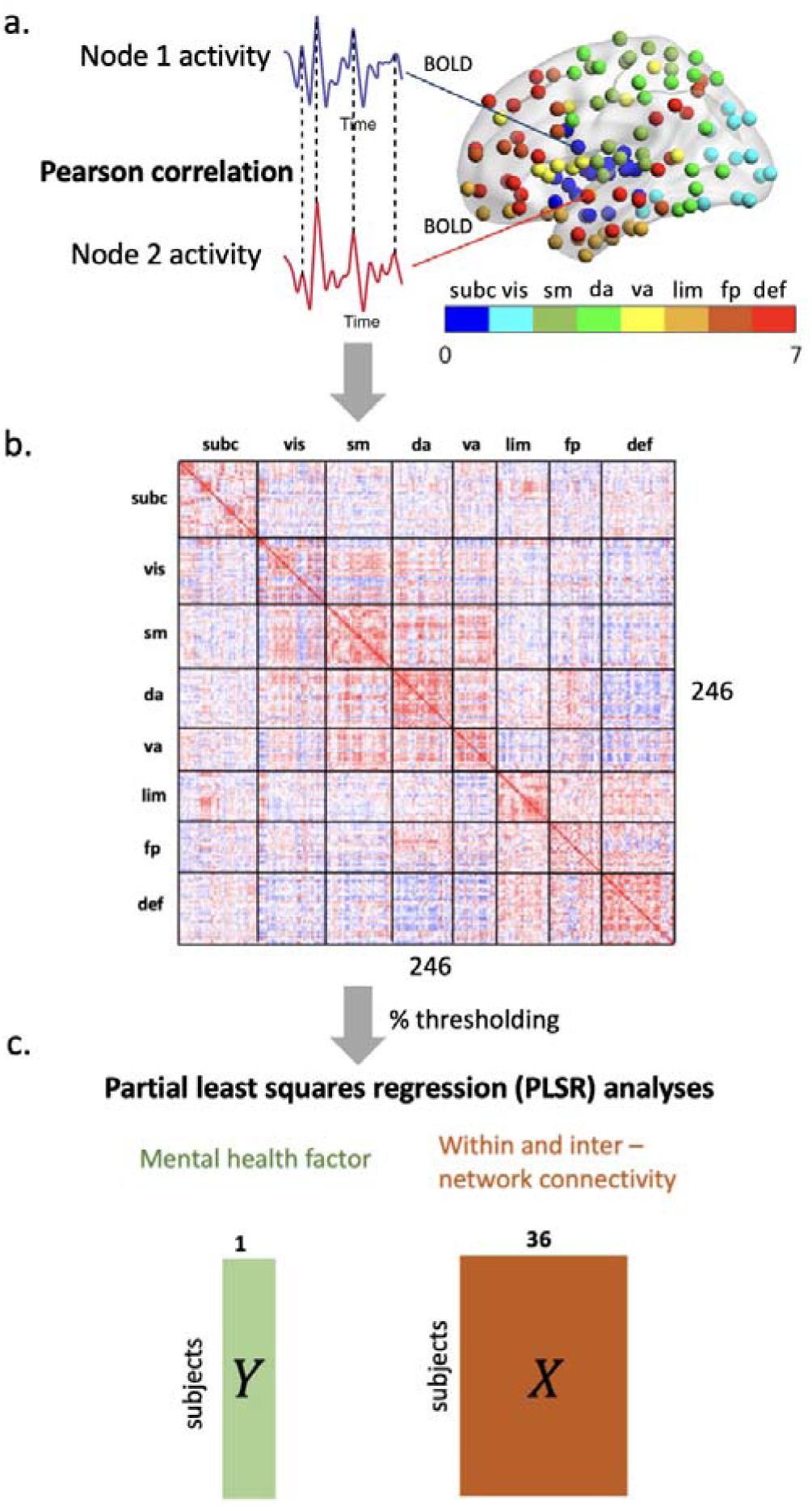
Schematic of the PLSR analysis workflow. a. The pre-processed rs-fMRI data is represented by blood oxygen-level dependent (BOLD) signal time series which were extracted from the 246 ROIs distributed across the cortex and subcortex. Nodes of the same colour belong to the same ICN as defined in (Yeo et al., 2011). b. Functional connectivity matrices (246 x 246) were constructed for all subjects in each of the two subsets (‘SDQ & Conners FC subset’ and ‘RCADS FC subset’), by calculating the Pearson correlation between the BOLD time-series of each pair of nodes (ROIs). c. Schematic representation of the two forms of the PLSR analyses performed in this study, as determined by the shape of the predictor matrix, X, which was calculated from the functional connectivity matrices of the subset of interest (‘SDQ & Conners FC subset’ or the ‘RCADS FC subset’). The matrix X (right panel) contains the functional connectivity within each of the 8 ICNs along the diagonal and the functional connectivity between each ICN pair (28 total pairs). Average regional functional connectivity for all subjects of a subset is represented by X, in the right panel. Both network-level and regional PLSR analyses were run six times, for each mental health dimension as the dependent variable.

Age, sex and motion (quantified as mean framewise displacement) were regressed from the edge-level functional connectivity data by fitting a linear model and calculating the residuals.

Intra- and inter-network connectivity was calculated as the average of edges within and between eight resting-state networks: subcortical, visual, somatomotor, dorsal-attention, ventral-attention, limbic, frontoparietal, default-mode network – DMN (Yeo et al., 2011).

#### Statistical analysis

Partial least squares regression (PLSR) was used to investigate the relationship between functional connectivity and the mental health dimensions. PLSR is a multivariate statistical method used to model relationships between two matrices (X and Y), by identifying components that explain the most covariance between the two matrices. PLSR was employed in the current study to investigate whether ICN-level functional connectivity could predict the variance in each derived mental health dimension (the dependent variable; Figure 1c). The predictor matrix consisted of within and between-network functional connectivity for the 8 ICNs (28 pairwise interactions), for all subjects. PLSR analyses were performed for each of the three externalising dimensions (externalising, neurodevelopmental and social maladjustment dimensions; N = 171) and for each of the p, internalising and specific internalising dimensions (N = 67) separately. Data from each mental health dimension was z-scored prior to the PLSR analysis. While the choice of deriving the externalising dimensions from the larger subset of participants with SDQ and Conners questionnaire data is advantageous for gaining statistical power for the hierarchical model of mental health and for the subsequent PLSR analyses, it makes direct comparisons between the results obtained with the externalising and internalising dimensions more difficult. As a robustness test, the PLSR analyses were also performed, in a later step, on the smaller subset of participants with data on all mental health dimensions and rs-fMRI data (see FigS4, FigS5 and TableS1 in Supplementary Results).

All PLSR analyses were performed in Scikit-learn version 0.20. To assess the significance of the PLSR components derived, 1000 permutations were performed on the dependent variable.

The PLSR analyses were also run with 1000 bootstraps of the dependent variable, to calculate variance of the PLSR x-loadings. The reported x-loadings were standardized by dividing the original loadings obtained from PLSR without permutations by the standard deviation of the x-loadings obtained in the bootstrapping procedure.

The significance of the PLSR components was determined by comparing the correlation between the x and y-scores with the correlation distribution between the x and y-scores from the 1000 permutations of the dependent variable. A significant threshold of alpha = 0.05 was used in the statistical tests to detect significant values. The obtained p-values from each of the six PLSR analyses with each mental health dimensions as the dependent variable were corrected for multiple comparisons (n = 6) using the false discovery rate (FDR). The methods employed in this study were previously pre-registered (Ianov Vitanov, 2025).

## Results

### Hierarchical model of mental health

Three latent dimensions could be maximally derived from the behavioural data from the SDQ, Conners and RCADS questionnaires using parallel analysis (Figure 2a). The number of dimensions retained (three) was determined by the number of eigenvalues from the real data that were larger than the eigenvalues from the simulated, random dataset. The simulated line in Figure 2a represents the upper limit of the 95% confidence interval around the simulated eigenvalues.

**Figure 2:**
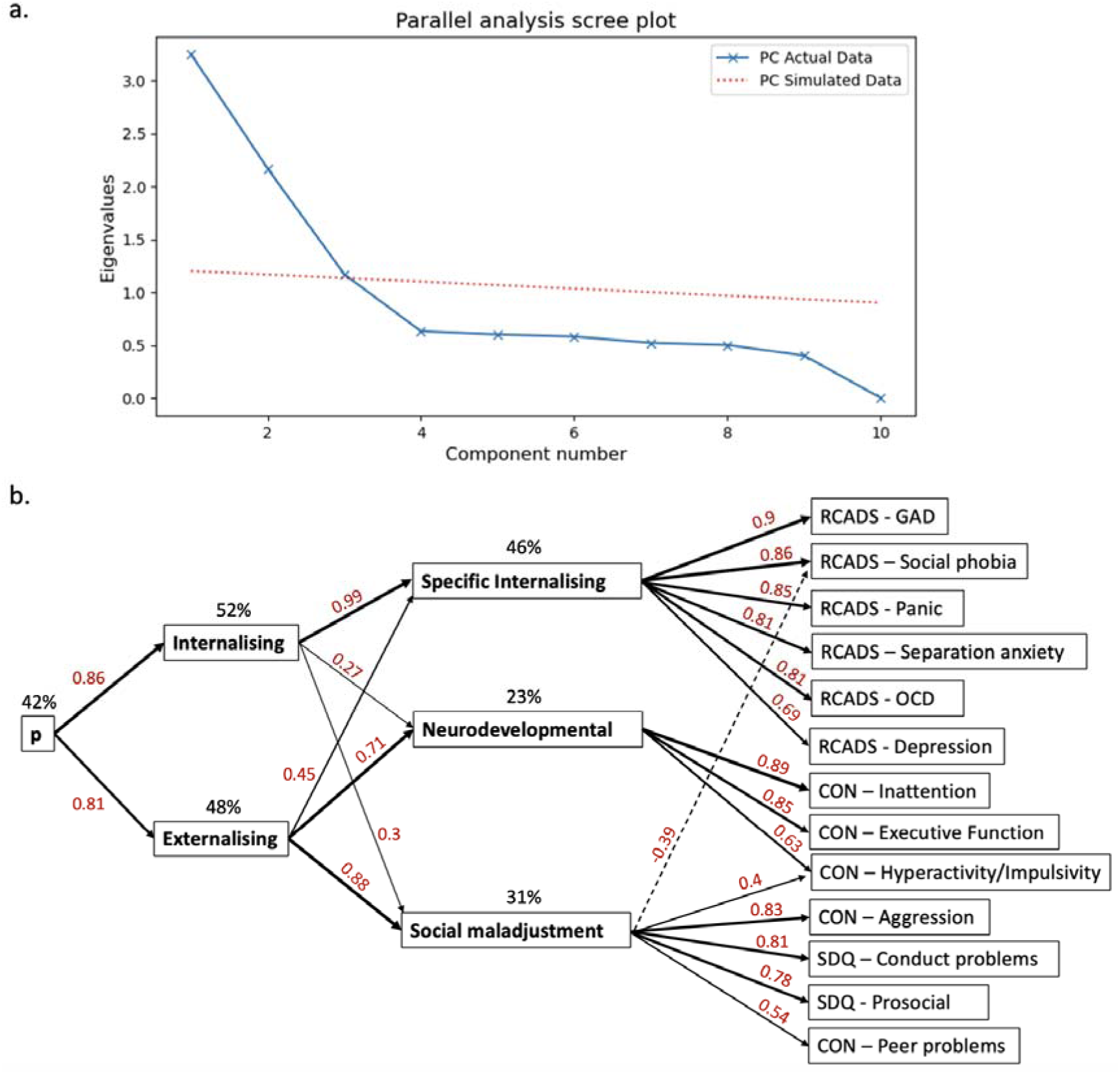
Results from the hierarchical model of mental health. a. Parallel analysis scree plot. b. Hierarchical structure of mental health in the CALM sample with all behavioural subscales of interest available (N = 378). The Spearman correlation coefficients, R between two adjacent mental health dimensions or between a mental health dimension and a questionnaire are shown in red, with the width of each arrow proportionate to the correlation coefficient. The dotted arrow indicates a negative correlation. Paths for absolute R < 0.3 are not shown. Figure format adapted from (Holmes et al., 2021). The variance in behavioural data explained by each mental health dimensions is shown above each dimensions. RCADS = Revised Child and Anxiety and Depression Scale (Parent Version). GAD = Generalised Anxiety Disorder. OCD = Obsessive Compulsive Disorder. CON = Conners-3 Parent Rating Scale Short Form. SDQ = Strengths and Difficulties Questionnaire.

At the highest level of the hierarchical model, a general p-factor captured a general predisposition for disrupted mental health (Figure 2b). At the second, lower level, the p-factor gave rise to the internalising and externalising dimensions, followed by the specific internalising, social maladjustment and the neurodevelopmental dimensions observed at the third level of the hierarchical model. The internalising dimensions fully predicted the specific internalising dimensions (0.99; p-value < 0.001). The externalising dimension was strongly associated with the neurodevelopmental dimensions (0.71; p-value < 0.001) and the social maladjustment dimensions (0.88; p-value < 0.001), respectively.

Pairwise correlations between the mental health dimensions of the paticipants with rs-fMRI data available are shown in Figure 3a. The correlations between each derived mental health dimension and the questionnaires scales are shown in Figure 3b. The p-factor was highly positively correlated with all included subscales, whereas the internalising and specific internalising dimensions showed strongest correlations with subscales indicating symptoms of anxiety and depression. The externalising dimension largely captured symptoms of social misconduct, aggression, as well as difficulties in attention, hyperactivity, altered executive function abilities, and, to a lower degree, depressive-like symptoms. The social maladjustment dimension was most strongly associated with subscales measuring disruptive social behaviour and to a lesser extent, hyperactivity, impulsivity and depression symptoms. In contrast, the neurodevelopmental dimension was most strongly correlated with patterns of behaviour that include inattention, hyperactivity/impulsivity and executive function difficulties. Overall, these findings closely mirrored those reported by Holmes et al. (2021), with identical mental health dimensions emerging and the relationships among them remaining consistent.

**Figure 3.**
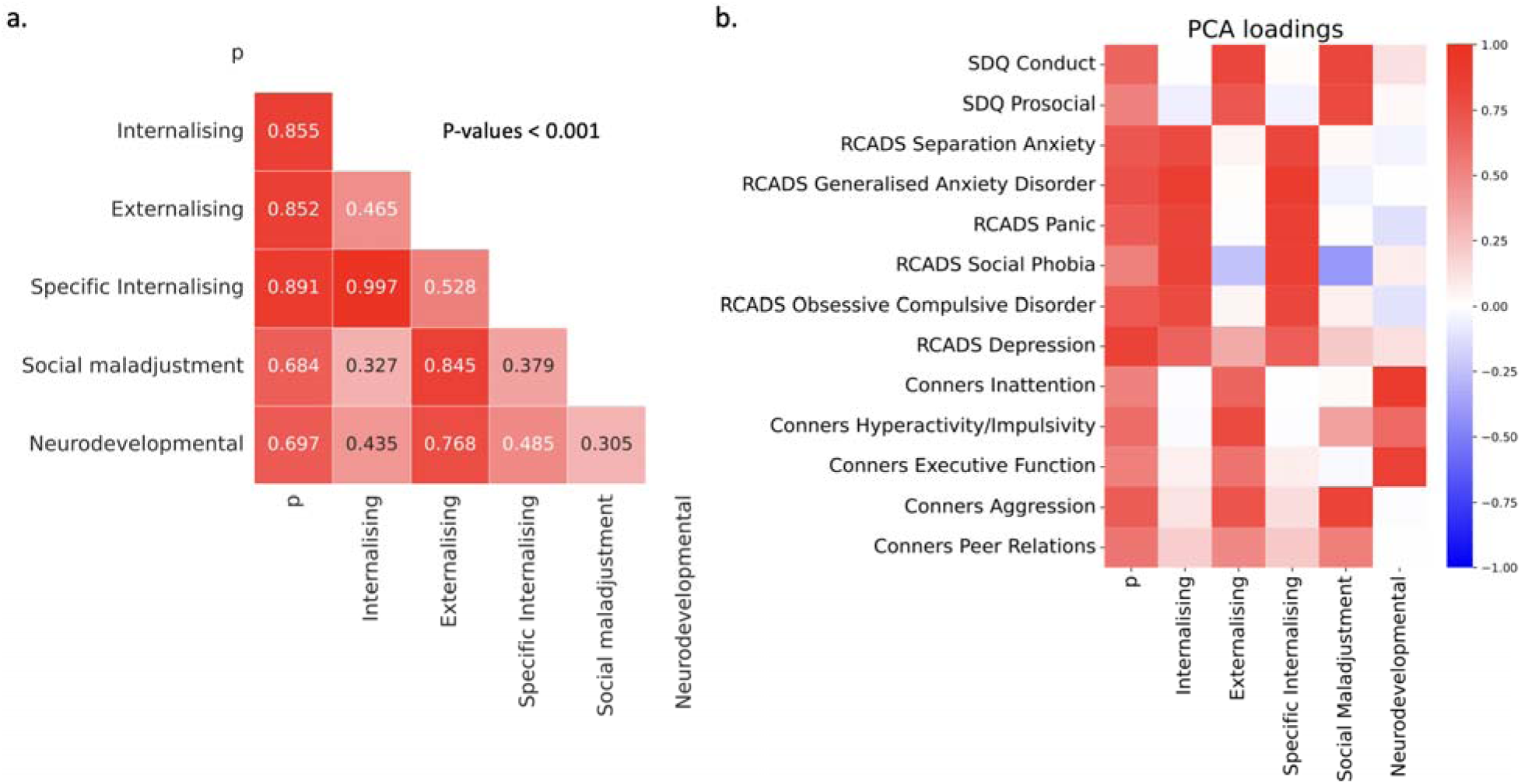
Dimensions correlations and loadings onto questionnaire subscales. a. Spearman correlation coefficients between all derived dimensions from the subsets of participants with rs-fMRI data. The correlation coefficient between the p-factor and the externalising dimensions denotes the strength of association between the p-factor scores and the externalising dimensions scores in the ‘RCADS FC subset’ (N = 67). To calculate the correlations between the externalising dimensions, the larger ‘SDQ and Conners FC subset’ was used (N = 171) . All correlations were significant (***p-values < 0.001). b. Dimensions loadings onto questionnaire subscales.

Associations between mental health dimensions and functional brain connectivity The Pearson correlations computed for each pair of nodal BOLD timeseries resulted in a 246 × 246 functional connectivity matrix for each subject. The edge values of the functional connectomes were centred at zero (FigS1) and motion effects on the rs-fMRI edge correlations were evaluated (FigS2). The residual motion effects were not statistically significant.

PLSR analyses revealed links between the mental health dimensions and network-level (within and between network) functional connectivity. Each PLSR analysis between each mental health dimensions and rs-FC resulted in only the first PLSR component being significant (p-value_FDR_ < 0.001), as resulting from the x-score and y-score correlations in the data compared to the correlations obtained when the behavioural variable was permuted. The correlation between the PLSR x and y-scores for each of the models are shown in Table 2. The latent components of FC explained 8 or 9% of variability in the respective mental health dimension that was included in a particular PLSR analysis. Although the correlation coefficients between functional connectivity and behavioural dimensions were statistically significant (R_S_ = .22–.34, all FDR-corrected p-values ≤ .008), the proportion of variance explained by the models was modest (8-9%) and did not survive FDR correction in most cases (Table 2). This indicates that while the latent connectivity patterns were reliably associated with behavioural measures, the overall explanatory power of the models was limited.

**Table 2:** Network-level PLSR results. Spearman correlation coefficient values between the PLSR-derived x-scores (representing a vector of latent FC within and between the 8 ICNs) and y-scores for each of the six network-level PLSR analyses performed, for each mental health dimension, and the associated FDR-corrected p-values are shown. The variance explained by the PLSR component from the corresponding network-level PLSR with each mental health dimensions and the associated uncorrected and FDR-corrected p-values are shown. P-values 0.05 are shown in bold.

| PLSR analysis | Correlation coefficient ($R_S$) | | Variance explained ($R_S^2$) | | |
| --- | --- | --- | --- | --- | --- |
| | $R_S$ value | p-value<br>(FDR-corrected) | $R_S^2$ value | p-value<br>(uncorrected) | p-value<br>(FDR-corrected) |
| p-factor | 0.22 | < **0.001** | 9% | 0.08 | 0.08 |
| Internalising | 0.34 | < **0.001** | 9% | 0.07 | 0.08 |
| Specific Internalising | 0.29 | < **0.001** | 8% | 0.08 | 0.08 |
| Externalising | 0.26 | < **0.001** | 9% | **0.02** | 0.1 |
| Neurodevelopmental | 0.27 | **0.008** | 8% | **0.05** | 0.1 |
| Social maladjustment | 0.24 | < **0.001** | 8% | **0.04** | 0.1 |

The correlation pattern between all the mental health dimensions scores (FigS3a) was largely recapitulated by the correlation pattern between the PLSR x-loadings, which reflected the latent components of brain connectivity derived from each PLSR analysis (FigS3b).

Next, the x-loadings onto the derived brain connectivity component from each PLSR analysis were investigated, as shown in Figure 4. We report these as a bootstrap ratio, by computing the mean loading across resamples divided by the bootstrap standard deviation. A positive x-loading reflects a positive correlation between a particular ICN feature and a behavioural variable, whereas a negative x-loading reflects a negative correlation. Most of the mental health dimensions were positively associated with increased functional connectivity within the DMN and between the DMN and the other ICNs (i.e. an increase in dimensions score, denoting heightened symptom severity, was positively associated with increased connectivity within network-level or between-network connectivity). The internalising dimensions showed small, positive associations with connectivity within the DMN and between the DMN and the frontoparietal network but had minimal association with connectivity between the DMN and the other ICNs. All six mental health dimensions were positively associated with functional connectivity within the frontoparietal network, as well as with connectivity between the frontoparietal network and the other ICNs. Qualitatively, these results suggest there may be both shared and specific patterns of network-level connectivity between the six mental health dimensions.

**Figure 4:**
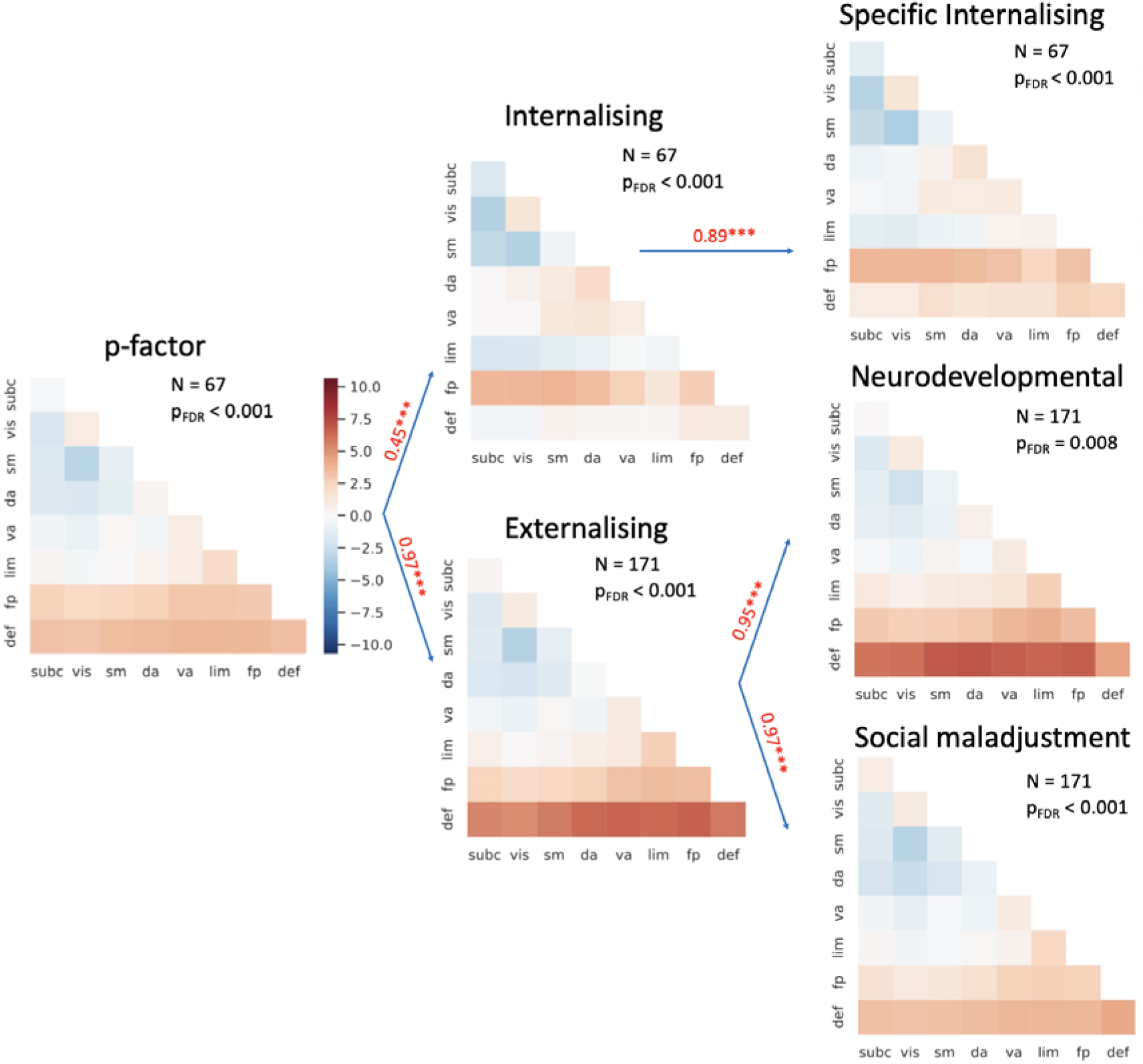
Network-level functional connectivity x-loadings from PLSR analyses performed at the network level, for each mental health dimensions. The x-loadings were computed as the mean loading across bootstrapping resamples divided by the standard deviation. The loadings are shown for each of the 8 included ICNs (abbreviations: subc – subcortical, vis – visual, sm – somatomotor, da – dorsal attention, va – ventral attention, lim – limbic, fp – frontoparietal, def – default-mode network) on the diagonal, and for each pairwise interaction between these 8 ICNs. The p, internalising and specific internalising dimensions scores were derived from the ‘RCADS FC subset’ (N = 67). The externalising, neurodevelopmental and social maladjustment dimensions were derived from the ‘SDQ & Conners subset’ (N = 171). The loadings which were positively correlated to each mental health dimensions are shown in red, with negative correlations shown in blue. The Spearman correlation coefficients between the network connectivity loadings for pairs of highly correlated mental health dimensions (R > 0.4) is shown on the arrows. The number of subjects for each PLSR analysis with the corresponding dimensions, as well as the FDR-corrected p-values from the correlations between the x and y-scores are shown (**p-value < 0.01; ***p-value < 0.001). The colorbar shown for the p-factor is valid for the other mental health dimensions shown in this figure.

We observed that higher functional integration (i.e. positive correlations between BOLD time-series) between the task-negative DMN and the task-positive executive networks (i.e. frontoparietal, dorsal attention, ventral attention) was positively correlated with all mental health dimensions. Greater integration between the DMN and subcortical, as well as sensory networks was positively associated with all mental health dimensions, except for the internalising dimensions, which showed weak negative correlations - possibly an artefact of the smaller sample size. All six mental health dimensions were also negatively associated with connectivity between the visual and somatomotor networks (i.e. an increase in dimensions score, indicating more severe symptoms, was negatively correlated with visual-somatomotor connectivity).

A visual inspection of Figure 4 suggested several differences in the connectivity patterns between some of the mental health dimensions. The connectivity between the dorsal-attention and ventral-attention networks was positively associated with the internalising and specific internalising dimensions, whereas the social maladjustment dimensions showed negative associations. The other dimensions did not show associations with this inter-network interaction. Increases in connectivity between the limbic network and the other ICNs were positively linked to the externalising and neurodevelopmental dimensions, whereas the internalising dimensions largely showed the opposite trend. These trends remain qualitative because the PLSR framework tests the significance of the latent brain-behaviour association rather than individual network loadings, and the current sample size limits reliable statistical comparison of loading patterns across dimensions.

## Discussion

Our results revealed relationships between functional brain connectivity and dimensions of mental health that captured internalising and externalising behaviours in a sample of youth at neurodevelopmental risk. Our hierarchical model of mental health symptoms distilled six symptom dimensions – spanning a broad p-factor down to specific internalising, social-maladjustment and neurodevelopmental facets. Strong coupling between the DMN and frontoparietal/attention systems emerged as a common neural correlate of psychopathology across these dimensions, mirroring prior transdiagnostic reports (ElliottllletZlal.,lll2018; XiallletZlal.,lll2018). This convergence highlights altered DMN-control network integration as a candidate neural signature of disrupted mental health in neurodevelopmental conditions.

Although these associations were statistically significant, the proportion of variance explained was modest (approximately 8–9%), indicating small but reliable effects. Functional connectivity therefore contributes to behavioural variation but accounts for only a limited proportion of individual differences. Such effect sizes are consistent with large-scale brain-wide association studies demonstrating that complex behavioural phenotypes are supported by distributed neural systems, each contributing modest effects rather than single, highly explanatory signatures (Marek *et al*., 2022).

At the level of specific dimensions, the general p-factor was associated with increased intrinsic connectivity within the DMN and frontoparietal networks, as well as enhanced coupling between these networks and other ICNs. These findings are consistent with previous reports demonstrating positive associations between general mental health risk and increased resting-state connectivity within the DMN and frontoparietal networks (Elliott *et al.,* 2018). Prior work has also identified heightened connectivity between the DMN and executive control networks (Xia *et al*., 2018), as well as between the DMN and subcortical networks (Hamilton *et al*., 2015).

Among p-factor-related connectivity patterns involving the DMN and frontoparietal networks, we observed positive associations with connectivity between the visual network and both the frontoparietal and the DMN. This pattern is consistent with prior findings (Elliott et al., 2018) and suggests that altered integration between visual processing regions and higher-order executive and task-negative systems is associated with general risk for poor mental health outcomes.

Similar ICN-behaviour associations were also evident across the externalising dimensions. The most pronounced effects emerged for the neurodevelopmental dimension, which was characterised by increased functional integration between the DMN and executive control networks, as well as between the DMN and subcortical and limbic systems.

Our results suggest that youth who showed executive-function deficits and inattention also exhibited reduced segregation (i.e., greater integration) between the DMN and task-positive control networks. This increased DMN–control coupling was the strongest functional-connectivity signature of the neurodevelopmental dimensions. This pattern contrasts with previous studies in healthy adults where greater DMN–control segregation predicted better attention (BarberllletZlal.,lll2015), working-memory accuracy (HampsonllletZlal.,lll2010) and lower impulsivity (InuggillletZlal.,lll2014). Those findings imply that keeping the DMN ‘offline’ while control networks are ‘online’ may be advantageous for goal-directed cognition. Yet our observation aligns with large-scale developmental work showing that stronger DMN–executive integration was linked to a range of mental-health difficulties, including externalising behaviour and mood lability (XiallletZlal.,lll2018). Taken together, the literature and our data point to a developmental shift: what is adaptive (DMN–control segregation) in mature cognition may fail to emerge – or even reverse – in youth at neurodevelopmental risk. Persistent DMN–control integration could therefore be a neural marker of the interplay between cognitive and affective dysregulation in this population.

The six mental health dimensions were negatively correlated with connectivity between the somatomotor and visual networks. Thus, higher scores for the mental health dimensions, denoting stronger mental health difficulties, was associated with lower connectivity between somatomotor and visual networks. This trend was previously found in an adult sample where increased p-factor scores were negatively linked to the FC between these ICNs (Elliott et al., 2018). Another study in neurotypical adults identified negative links between somatomotor-visual connectivity and impulsivity levels, which might highlight a disrupted process in neurodevelopmental symptoms in which perceptual information from sensory cortices is less integrated with behavioural control programs (Herman et al., 2020). The current results indicated, at a qualitative level, that the strongest association existed between reduced visual-somatomotor connectivity and the social maladjustment dimensions. Moreover, the negative associations between visual-somatomotor network connectivity and the internalising as well as the externalising dimensions highlight the possibility that internalising mental health difficulties could co-occur and might be underpinned by some similar connectivity patterns with impulsivity and other symptoms that are typical for externalising disruptive behaviour. This would be in line with previous studies showing comorbidities between internalising symptoms and those of inattention, poor executive function and conduct problems, which underlie the externalising dimensions (Doering et al., 2021; Kerns et al., 2014; Bauermeister et al., 2007).

Somatomotor network connectivity has also been previously related to transdiagnostic dimensions of mental health displaying altered connectivity within this large-scale network and between the somatomotor network and subcortical, as well as cortical executive networks (Kebets et al., 2019), suggesting the central role of the somatomotor network in information processing relevant to behavioural dysregulation. Future studies should investigate the directionality of these processes in neurodevelopmental risk.

Finally, dissociable patterns of connectivity were also identified between the mental health dimensions. Analyses conducted in the smaller subsample (N = 67; FigS5) suggest weaker, positive associations between the internalising dimensions and functional connectivity within the DMN, as well as between the DMN and other ICNs, relative to the p-factor and externalising dimensions. However, future studies should validate these findings in larger samples with a balanced number of participants across mental health dimensions. Previous studies identified positive links between increased DMN connectivity and mind wandering in depression (Rosenbaum et al., 2017), as well as with symptom severity in recurrent major depressive disorder (Yan et al., 2019), highlighting the key implication of this network in internalising symptoms. In line with previous research (Xia et al., 2018), the derived internalising dimensions in our model also appeared to be weakly positively correlated with connectivity between the dorsal-attention and the ventral attention networks (FigS5), but future validations in larger sample sizes are required to clarify this association.

### Study limitations and potential future directions

There are several limitations of this study which should be considered in the light of the results and their interpretations. One of these is the smaller subset of participants with data on the RCADS questionnaire. Similar connectivity patterns were obtained when the ‘RCADS FC subset’ (N = 67) was used to test for associations between each mental health dimensions and functional connectivity (see FigS5). However, this analysis comes with a large reduction in statistical power and thus the difference in sample size makes it difficult to robustly compare the relationships between connectivity and different mental health dimensions. This work should be validated in larger datasets where behavioural and functional connectivity data are available across the sample. Another limitation of the study is our focus on behavioural and MRI data from a single timepoint (i.e. cross-sectional data). In future, it will be essential to collect behavioural and MRI data at different timepoints in childhood and adolescence to assess the temporal unfolding and potential causal trends in mental health divergence, as well as to capture greater effect sizes and statistical power (Marek et al., 2022). Lastly, going forwards, sampling endeavours should consider subscales that capture a wider range of behaviours to better characterise mental health. The six derived mental health dimensions from the available behavioural questionnaires were constrained by the breadth of the available subscales. Other studies which derived mental health dimensions in different samples obtained other dimensions such as psychosis (Xia et al., 2018), somatoform and detachment (Karcher et al., 2021), suggesting the existence of additional relevant dimensions. Designing more varied and granular mental health subscales that capture a larger number of behavioural traits as well employing a more holistic sampling process that would include physiological, environmental and genetic data will be essential to obtain a better understanding of the complexity of the mental health landscape.

In conclusion, we demonstrate that a hierarchically organised, six-dimensions model of disrupted mental health in neurodevelopmentally at-risk adolescents is underpinned by two robust signatures of resting-state functional brain connectivity. First, progressively stronger coupling of the DMN with frontoparietal and attention systems tracked symptom severity from the transdiagnostic p-factor down to specific mental health dimensions – implicating DMN-control hyper-integration as a unifying circuit marker of disorder risk. Second, diminished connectivity between visual and somatomotor networks, most pronounced for the social-maladjustment dimension, delineated a complementary motor–sensory pathway of dysfunction. These convergent motifs offer a succinct neural fingerprint of cognitive-affective dysregulation and nominate DMN-control integration as a potential biomarker for transdiagnostic mental-health vulnerability in youth.

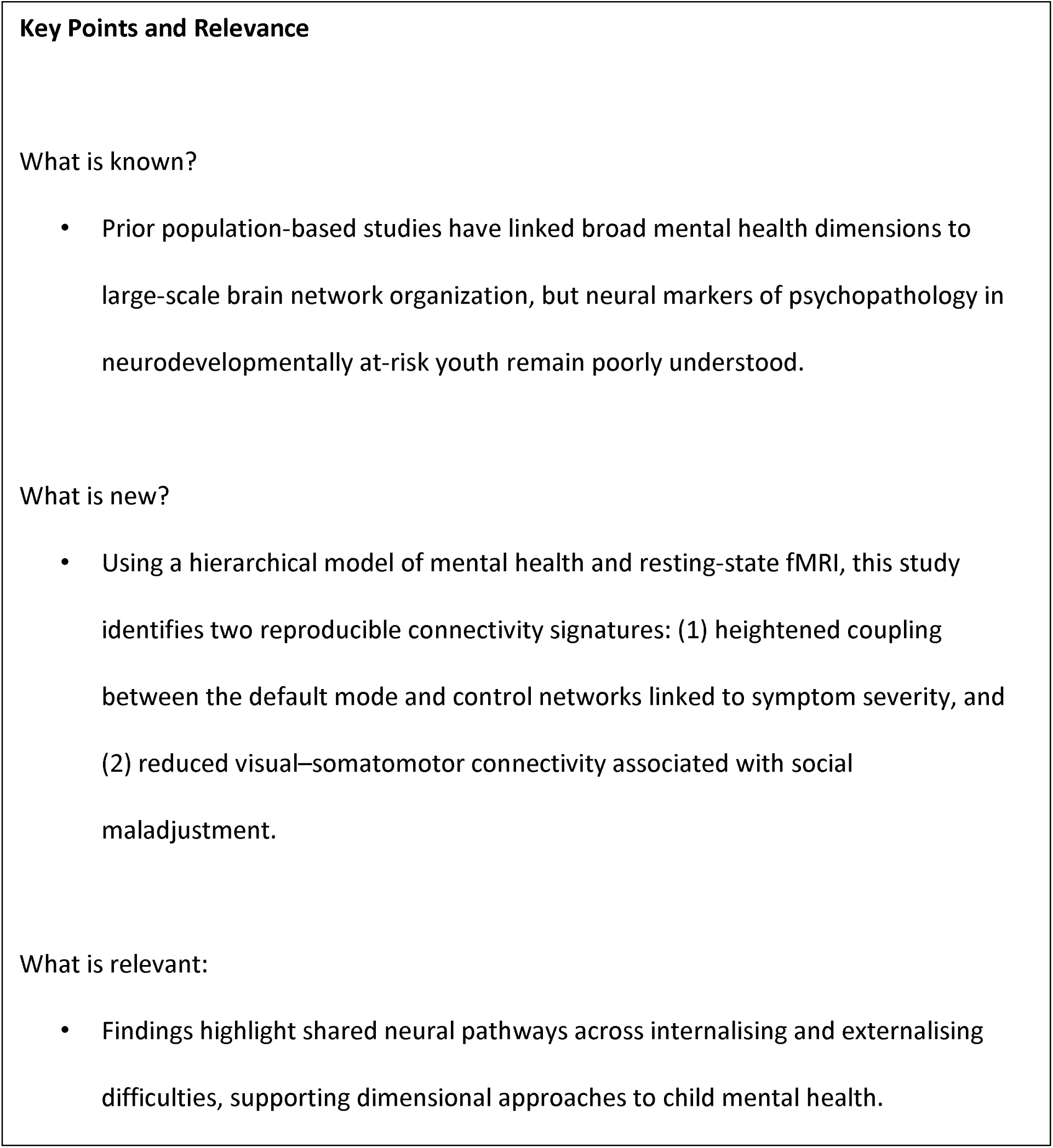

## ACKNOWLEDGEMENTS

Data collection and sharing were conducted by the Centre for Attention, Learning and Memory (CALM), funded by the UK Medical Research Council and the University of Cambridge. Data used in this study were accessed via the CALM resource – https://calm.mrc-cbu.cam.ac.uk/. The study protocol is reported in Holmes et al. (2019).

The authors were supported by the Medical Research Council program grant MC-A0606-5PQ41. We would like to thank all members of the CALM Team for their help with recruitment, data collection, and data management, as well as all of the children and parents for their participation in the study. The Centre for Attention Learning and Memory (CALM) research clinic is based at the Medical Research Council (MRC) Cognition and Brain Sciences Unit, University of Cambridge, funded by the MRC through UKRI. The lead investigators are Duncan Astle, Kate Baker, Susan Gathercole, Joni Holmes, Rogier Kievit and Tom Manly. Data collection is assisted by a team of researchers and PhD students that includes Danyal Akarca, Joe Bathelt, Marc Bennett, Madalena Bettencourt, Giacomo Bignardi, Sarah Bishop, Erica Bottacin, Lara Bridge, Diandra Brkic, Annie Bryant, Sally Butterfield, Elizabeth Byrne, Gemma Crickmore, Edwin Dalmaijer, Fánchea Daly, Tina Emery, Laura Forde, Grace Franckel, Delia Furhmann, Andrew Gadie, Sara Gharooni, Jacalyn Guy, Erin Hawkins, Rebeca Ianov-Vitanov, Christian Iordanov, Agnieszka Jaroslawska, Sara Joeghan, Amy Johnson, Jonathan Jones, Silvana Mareva, Jessica Martin, Alicja Monaghan, Elise Ng-Cordell, Sinead O’Brien, Cliodhna O’Leary, Joseph Rennie, Andrea Santangelo, Ivan Simpson-Kent, Roma Siugzdaite, Tess Smith, Stepheni Uh, Maria Vedechkina, Francesca Woolgar, Natalia Zdorovtsova, Mengya Zhang. The authors wish to thank the many professionals working in children’s services in the South-East and East of England for their support, and to the children and their families for giving up their time to visit the clinic. We are also very appreciative of the radiographers at the MRC Cognition and Brain Sciences Unit for their expertise and continued support in delivering outstanding paediatric MRI scanning.

## ETHICS APPROVAL STATEMENT

The original CALM study received ethical approval on 27/08/2013 from the National Health Service (Approval Reference Number: 13/EE/0157) and was conducted in accordance with the Declaration of Helsinki. This secondary analysis was approved by the CALM Management Committee.

## DATA AVAILABILITY STATEMENT

Data were obtained through the CALM: Open portal. Anonymised clinical, cognitive, structural, and functional MRI data are available to researchers at academic and clinical institutions worldwide for non-commercial use, subject to application and approval by the CALM Management Committee and agreement to its ethical terms. Further information is available via the CALM portal or by contacting.

## PATIENT CONSENT STATEMENT

Written informed consent was obtained from all participants and their parents/carers as part of the original CALM study (Holmes et al., 2019). The present secondary analysis used anonymised data accessed through the CALM: Open portal.

## PERMISSION TO REPRODUCE MATERIAL FROM OTHER SOURCES

No material requiring permission from other sources has been reproduced.

## CLINICAL TRIAL REGISTRATION

Not applicable.

## Abbreviations

BOLD: Blood oxygen level–dependent
CALM: Centre for Attention, Learning and Memory
DMN: Default mode network
fMRI: Functional magnetic resonance imaging
FPN: Frontoparietal network
ICN: Intrinsic connectivity network
PCA: Principal component analysis
PLSR: Partial least squares regression
QC: Quality control
RCADS: Revised Children’s Anxiety and Depression Scale
RDoC: Research Domain Criteria
RSFC: Resting-state functional connectivity
SDQ: Strengths and Difficulties Questionnaire

## References

1. Agnafors, S., Barmark, M. and Sydsjö, G. (2021). Mental health and academic performance: a study on selection and causation effects from childhood to early adulthood. Social psychiatry and psychiatric epidemiology 56(5), 857–866.

2. American Psychiatric Association. (2022). Diagnostic and statistical manual of mental disorders (5th ed., text rev.)

3. Astle, D. E., Bathelt, J., CALM Team, & Holmes, J. (2019). Remapping the cognitive and neural profiles of children who struggle at school. Developmental Science, 22(1), e12747.

4. Barber, A. D., Jacobson, L. A., Wexler, J. L., Nebel, M. B., Caffo, B. S., Pekar, J. J., & Mostofsky, S. H. (2015). Connectivity supporting attention in children with attention deficit hyperactivity disorder. NeuroImage: Clinical, 7, 68–81.

5. Bathelt, J., Holmes, J., Astle, D. E., Gathercole, S., Manly, T., & Kievit, R. (2018). Data-driven subtyping of executive function–related behavioral problems in children. Journal of the American Academy of Child & Adolescent Psychiatry, 57(4), 252–262.

6. Bauermeister, J. J., Shrout, P. E., Ramírez, R., Bravo, M., Alegría, M., Martínez-Taboas, A., Chávez, L., Rubio-Stipec, M., García, P., & Ribera, J. C. (2007). ADHD correlates, comorbidity, and impairment in community and treated samples of children and adolescents. Journal of Abnormal Child Psychology, 35(6), 883–898.

7. Behzadi, Y., Restom, K., Liau, J., & Liu, T. T. (2007). A component based noise correction method (CompCor) for BOLD and perfusion based fMRI. NeuroImage, 37(1), 90–101.

8. Bethlehem, R. A. I., Seidlitz, J., White, S. R., Vogel, J. W., Anderson, K. M., Adamson, C., Adler, S., Alexopoulos, G. S., Anagnostou, E., Areces-Gonzalez, A., et al. (2022). Brain charts for the human lifespan. Nature, 604(7906), 525–533.

9. Cao, M., Shu, N., Cao, Q., Wang, Y., & He, Y. (2014). Imaging functional and structural brain connectomics in attention-deficit/hyperactivity disorder. Molecular Neurobiology, 50(3), 1111–1123.

10. Casey, B. J., Cannonier, T., Conley, M. I., Cohen, A. O., Barch, D. M., Heitzeg, M. M., Soules, M. E., Teslovich, T., Dellarco, D. V., Garavan, H., Orr, C. A., Wager, T. D., Banich, M. T., Speer, N. K., Sutherland, M. T., Riedel, M. C., Dick, A. S., Bjork, J. M., Thomas, K. M., Chaarani, B., … ABCD Imaging Acquisition Workgroup (2018). The Adolescent Brain Cognitive Development (ABCD) study: Imaging acquisition across 21 sites. Developmental cognitive neuroscience, 32, 43–54.

11. Caspi, A., Houts, R. M., Belsky, D. W., Goldman-Mellor, S. J., Harrington, H., Israel, S., Meier, M. H., Ramrakha, S., Shalev, I., & Poulton, R. (2014). The p dimensions: One general psychopathology dimensions in the structure of psychiatric disorders? Clinical Psychological Science, 2(2), 119–137.

12. Chorpita, B. F., Yim, L., Moffitt, C., Umemoto, L. A., & Francis, S. E. (2000). Assessment of symptoms of DSM-IV anxiety and depression in children: A revised child anxiety and depression scale. Behaviour Research and Therapy, 38(8), 835–855.

13. Conners, C. K. (2008). Conners 3–Parent Short Form. Multi-Health Systems.

14. Cox, R. W. (1996). AFNI: Software for analysis and visualization of functional magnetic resonance neuroimages. Computers and Biomedical Research, 29(3), 162–173.

15. Cuthbert, B. N. (2014). The RDoC framework: Facilitating transition from ICD/DSM to dimensional approaches that integrate neuroscience and psychopathology. World Psychiatry, 13(1), 28–35.

16. Cuthbert, B. N. (2022). Research domain criteria (RDoC): Progress and potential. Current Directions in Psychological Science, 31(2), 107–114.

17. Doering, S., Lichtenstein, P., Gillberg, C., Kuja-Halkola, R., & Lundström, S. (2021). Internalizing and neurodevelopmental problems in young people: Educational outcomes in a large population-based cohort of twins. Psychiatry Research, 298, 113794.

18. Donati, G., Meaburn, E., & Dumontheil, I. (2021). Internalising and externalising in early adolescence predict later executive function, not the other way around: A cross-lagged panel analysis. Cognition and Emotion, 35(5), 986–998.

19. Elliott, M. L., Romer, A., Knodt, A. R., & Hariri, A. R. (2018). A connectome-wide functional signature of transdiagnostic risk for mental illness. Biological Psychiatry, 84(6), 452–459.

20. Esteban, O., Markiewicz, C. J., Blair, R. W., Moodie, C. A., Isik, A. I., Erramuzpe, A., Kent, J. D., Goncalves, M., DuPre, E., & Snyder, M. (2019). fMRIPrep: A robust preprocessing pipeline for functional MRI. Nature Methods, 16(1), 111–116.

21. Fan, L., Li, H., Zhuo, J., Zhang, Y., Wang, J., Chen, L., Yang, Z., Chu, C., Xie, S., & Laird, A. R. (2016). The human brainnetome atlas: A new brain atlas based on connectional architecture. Cerebral Cortex, 26(8), 3508–3526.

22. Goldberg, L. R. (2006). Doing it all bass-ackwards: The development of hierarchical dimensions structures from the top down. Journal of Research in Personality, 40(4), 347–358.

23. Goodman, A., & Goodman, R. (2009). Strengths and Difficulties Questionnaire as a dimensional measure of child mental health. Journal of the American Academy of Child & Adolescent Psychiatry, 48(4), 400–403.

24. Grydeland, H., Vértes, P. E., Váša, F., Romero-Garcia, R., Whitaker, K., Alexander-Bloch, A. F., Bjørnerud, A., Patel, A. X., Sederevičius, D., & Tamnes, C. K. (2019). Waves of maturation and senescence in micro-structural MRI markers of human cortical myelination over the lifespan. Cerebral Cortex, 29(3), 1369–1381.

25. Hamilton, J. P., Farmer, M., Fogelman, P., & Gotlib, I. H. (2015). Depressive rumination, the default-mode network, and the dark matter of clinical neuroscience. Biological Psychiatry, 78(4), 224–230.

26. Hampson, M., Driesen, N., Roth, J. K., Gore, J. C., & Constable, R. T. (2010). Functional connectivity between task-positive and task-negative brain areas and its relation to working memory performance. Magnetic Resonance Imaging, 28(8), 1051–1057.

27. Hansen, B. H., Oerbeck, B., Skirbekk, B., Petrovski, B. É., & Kristensen, H. (2018). Neurodevelopmental disorders: Prevalence and comorbidity in children referred to mental health services. Nordic Journal of Psychiatry, 72(4), 285–291.

28. Herman, A. M., Critchley, H. D., & Duka, T. (2020). Trait impulsivity associated with altered resting-state functional connectivity within the somatomotor network. Frontiers in Behavioral Neuroscience, 14, 111.

29. Holmes, J., Bryant, A., & Gathercole, S. E. (2019). Protocol for a transdiagnostic study of children with problems of attention, learning and memory (CALM). BMC Pediatrics, 19(1), 1–11.

30. Holmes, J., Mareva, S., Bennett, M. P., Black, M. J., & Guy, J. (2021). Higher-order dimensions of psychopathology in a neurodevelopmental transdiagnostic sample. Journal of Abnormal Psychology, 130(8), 909–920.

31. Horn, J. L. (1965). A rationale and test for the number of dimensions in dimensions analysis. Psychometrika, 30(2), 179–185.

32. Ianov Vitanov, R. (2025). Characterising links between brain connectivity and mental health in a transdiagnostic sample of struggling learners. OSF. 10.17605/OSF.IO/64ZSK .

33. Insel, T. R., Cuthbert, B. N., Garvey, M., Heinssen, R., Pine, D. S., Quinn, K., Sanislow, C., & Wang, P. (2010). Research Domain Criteria (RDoC): Toward a new classification framework for research on mental disorders. American Journal of Psychiatry, 167(7), 748–751.

34. Inuggi, A., Sanz-Arigita, E., González-Salinas, C., Valero-García, A. V., García-Santos, J. M., & Fuentes, L. J. (2014). Brain functional connectivity changes in children that differ in impulsivity temperamental trait. Frontiers in Behavioral Neuroscience, 8, 156.

35. Jones, J. S., Astle, D. E., CALM Team, & Holmes, J. (2021). A transdiagnostic data-driven study of children’s behaviour and the functional connectome. Developmental Cognitive Neuroscience, 52, 101027.

36. Jones, J. S., CALM Team, & Astle, D. E. (2022). Segregation and integration of the functional connectome in neurodevelopmentally ‘at risk’ children. Developmental Science, 25(3), e13209.

37. Kapadia, M., Desai, M., & Parikh, R. (2020). Fractures in the framework: Limitations of classification systems in psychiatry. Dialogues in Clinical Neuroscience, 22(1), 17–26.

38. Karcher, N. R., Michelini, G., Kotov, R., & Barch, D. M. (2021). Associations between resting-state functional connectivity and a hierarchical dimensional structure of psychopathology in middle childhood. Biological Psychiatry: Cognitive Neuroscience and Neuroimaging, 6(5), 508–517.

39. Kebets, V., Holmes, A. J., Orban, C., Tang, S., Li, J., Sun, N., Kong, R., Poldrack, R. A., & Yeo, B. T. (2019). Somatosensory-motor dysconnectivity spans multiple transdiagnostic dimensions of psychopathology. Biological Psychiatry, 86(10), 779–791.

40. Kerns, C. M., Kendall, P. C., Berry, L., Souders, M. C., Franklin, M. E., Schultz, R. T., Miller, J., & Herrington, J. (2014). Traditional and atypical presentations of anxiety in youth with autism spectrum disorder. Journal of Autism and Developmental Disorders, 44(11), 2851–2861.

41. Kotov, R., Cicero, D. C., Conway, C. C., DeYoung, C. G., Dombrovski, A. Y., Eaton, N. R., First, M. B., Forbes, M. K., Hyman, S. E., & Jonas, K. G. (2022). The hierarchical taxonomy of psychopathology (HiTOP) in psychiatric practice and research. Psychological Medicine, 52(9), 1666–1678.

42. Kotov, R., Krueger, R. F., Watson, D., Achenbach, T. M., Althoff, R. R., Bagby, R. M., Brown, T. A., Carpenter, W. T., Caspi, A., & Clark, L. A. (2017). The hierarchical taxonomy of psychopathology (HiTOP): A dimensional alternative to traditional nosologies. Journal of Abnormal Psychology, 126(4), 454–477.

43. Lahey, B. B., Moore, T. M., Kaczkurkin, A. N., & Zald, D. H. (2021). Hierarchical models of psychopathology: Empirical support, implications, and remaining issues. World Psychiatry, 20(1), 57–63.

44. Marek, S., Tervo-Clemmens, B., Calabro, F. J., Montez, D. F., Kay, B. P., Hatoum, A. S., Donohue, M. R., Foran, W., Miller, R. L., & Hendrickson, T. J. (2022). Reproducible brain-wide association studies require thousands of individuals. Nature, 603(7902), 654–660.

45. Milienos, F. S., Rentzios, C., Catrysse, L., Gijbels, D., Mastrokoukou, S., Longobardi, C., & Karagiannopoulou, E. (2021). The contribution of learning and mental health variables in first-year students’ profiles. Frontiers in Psychology, 12, 627118.

46. Monaghan, A., Misic, B., Shafiei, G., Tsvetanov, K. A., Astle, D. E., & Bethlehem, R. A. (2026). Integrating Functional Transdiagnostic Dimensions of Psychopathology with Cortical Organization. bioRxiv, 2026-02.

47. Morris-Rosendahl, D. J., & Crocq, M.-A. (2022). Neurodevelopmental disorders—The history and future of a diagnostic concept. Dialogues in Clinical Neuroscience, 24(3), 295–308.

48. Munir, K. M. (2016). The co-occurrence of mental disorders in children and adolescents with intellectual disability/intellectual developmental disorder. Current Opinion in Psychiatry, 29(2), 95–102.

49. Murphy, J. M., Guzmán, J., McCarthy, A. E., Squicciarini, A. M., George, M., Canenguez, K. M., Dunn, E. C., Baer, L., Simonsohn, A., & Smoller, J. W. (2015). Mental health predicts better academic outcomes: A longitudinal study of elementary school students in Chile. Child Psychiatry & Human Development, 46(2), 245–256.

50. Nomi, J. S., & Uddin, L. Q. (2015). Developmental changes in large-scale network connectivity in autism. NeuroImage: Clinical, 7, 732–741.

51. Power, J. D., Barnes, K. A., Snyder, A. Z., Schlaggar, B. L., & Petersen, S. E. (2012). Spurious but systematic correlations in functional connectivity MRI networks arise from subject motion. NeuroImage, 59(3), 2142–2154.

52. Rosenbaum, D., Haipt, A., Fuhr, K., Haeussinger, F. B., Metzger, F. G., Nuerk, H.-C., Fallgatter, A. J., Batra, A., & Ehlis, A.-C. (2017). Aberrant functional connectivity in depression as an index of state and trait rumination. Scientific Reports, 7(1), 2174.

53. Satterthwaite, T. D., Elliott, M. A., Ruparel, K., Loughead, J., Prabhakaran, K., Calkins, M. E., Hopson, R., Jackson, C., Keefe, J., & Riley, M. (2014). Neuroimaging of the Philadelphia Neurodevelopmental Cohort. NeuroImage, 86, 544–553.

54. Satterthwaite, T. D., Wolf, D. H., Loughead, J., Ruparel, K., Elliott, M. A., Hakonarson, H., Gur, R. C., & Gur, R. E. (2012). Impact of in-scanner head motion on multiple measures of functional connectivity: Relevance for studies of neurodevelopment in youth. NeuroImage, 60(1), 623–632.

55. Sripada, C. S., Kessler, D., & Angstadt, M. (2014). Lag in maturation of the brain’s intrinsic functional architecture in attention-deficit/hyperactivity disorder. Proceedings of the National Academy of Sciences, 111(39), 14259–14264.

56. Supekar, K., Musen, M., & Menon, V. (2009). Development of large-scale functional brain networks in children. PLoS Biology, 7(7), e1000157.

57. van den Heuvel, M. P., Stam, C. J., Boersma, M., & Hulshoff Pol, H. E. (2008). Small-world and scale-free organization of voxel-based resting-state functional connectivity in the human brain. NeuroImage, 43(3), 528–539.

58. Wakefield, J. C. (2015). DSM-5, psychiatric epidemiology and the false-positives problem. Epidemiology and Psychiatric Sciences, 24(3), 188–196.

59. Xia, C. H., Ma, Z., Ciric, R., Gu, S., Betzel, R. F., Kaczkurkin, A. N., Calkins, M. E., Cook, P. A., García de la Garza, A., & Vandekar, S. N. (2018). Linked dimensions of psychopathology and connectivity in functional brain networks. Nature Communications, 9(1), 3003.

60. Yan, C.-G., Chen, X., Li, L., Castellanos, F. X., Bai, T.-J., Bo, Q.-J., Cao, J., Chen, G.-M., Chen, N.-X., & Chen, W. (2019). Reduced default mode network functional connectivity in patients with recurrent major depressive disorder. Proceedings of the National Academy of Sciences, 116(18), 9078–9083.

61. Yeo, B. T. T., Krienen, F. M., Sepulcre, J., Sabuncu, M. R., Lashkari, D., Hollinshead, M., Roffman, J. L., Smoller, J. W., Zöllei, L., & Polimeni, J. R. (2011). The organization of the human cerebral cortex estimated by intrinsic functional connectivity. Journal of Neurophysiology, 106(3), 1125–1165.

