## Supplemental Information for "Functional connectivity correlates of the hierarchical p-factor model in youth at neurodevelopmental risk"

**Supplementary Methods**

**Resting-state functional MRI data**

The fMRI data were originally pre-processed by Jones *et al*. (2021), and full details of the pre-processing steps are provided there. Briefly, the available fMRI data was minimally pre-processed in fMRIPrep version 1.5.0 (Esteban *et al*., 2019), which implements slice-timing correction, rigid-body realignment, boundary-based co-registration to the structural T1, segmentation, and normalisation to the MNI template. The data was used without smoothing. The fMRI data was denoised using the fmridenoise package in Python. The most effective confound regression procedure included a band-pass filter between 0.01 and 0.1 Hz, 24 head motion parameters (six rigid body realignment parameters, their squares, their derivatives, and their squared derivatives), 10 aCompCor components from the WM and CSF signal (Behzadi *et al.*, 2007), linear and quadratic trends, and motion spikes (framewise displacement > 0.5 mm; (Power *et al*., 2012). Simultaneous confound regression was performed in the Nipype (version 1.2.0) implementation of AFNI’s 3dTproject (Cox, 1996). Children were first excluded for high average motion (mean framewise displacement > 0.5 mm, n = 93) and then for a large number of motion spikes (> 20% spikes, n = 18), where few temporal degrees of freedom would have remained. The final functional connectome sample included 241 children.

**Supplementary Results**

Functional connectivity results

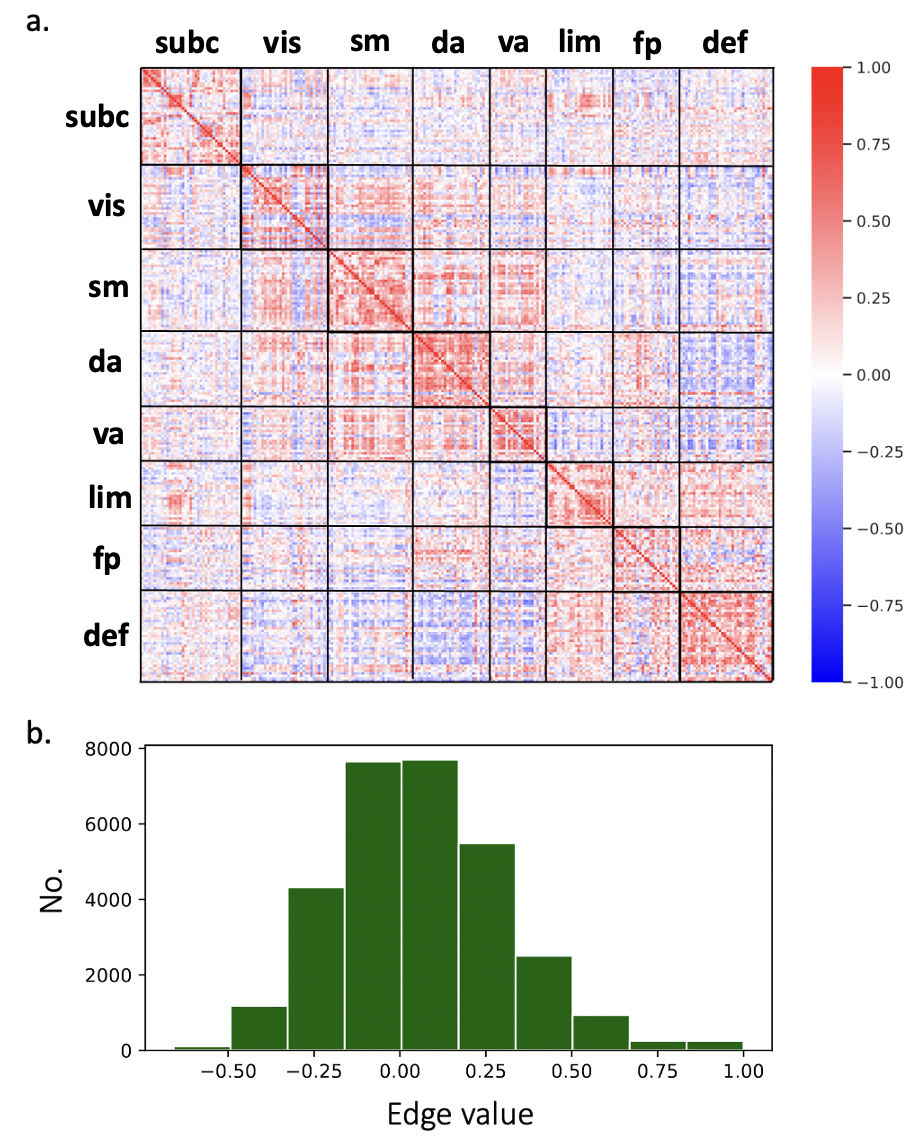

**FigS1**: a. Unthresholded functional connectivity matrix, from one of the subjects, parcellated according to the seven Yeo networks: subcortical (subc), visual (vis), somatomotor (sm), dorsal- attention (da), ventral-attention (va), limbic (lim), frontoparietal (fp), default-mode network (def). b. Edge value distribution from the unthresholded functional connectivity matrix of one subject, shown as an example.

The effects of motion on the rs-fMRI edge correlations were investigated to exclude the

possibility that in-scanner head motion, which is a confound, did not drive the rs-fMRI

results. The mean framewise displacement (Mean FD) was used to quantify motion in

the sample (FigS2a). Mean FD did not show a significant association with the mean

functional connectivity (Mean FC) ($R$ = -0.12; *p-value* = 0.11; FigS2b). The degree

to which edge length (the Euclidean distance between a node pair) modulates the effect

of motion on edge-wise connectivity (Satterthwaite *et al*., 2012) was also assessed. This

was achieved by correlating each edge from the unthresholded functional connectivity

matrices to mean framewise displacement (Mean FD) across subjects, and the correlation

coefficient obtained was then correlated with edge length. The influence of edge length on

the correlation between motion and edge-level functional connectivity was small and not

significant (R=0.007; *p-value* = 0.18; FigS2c).

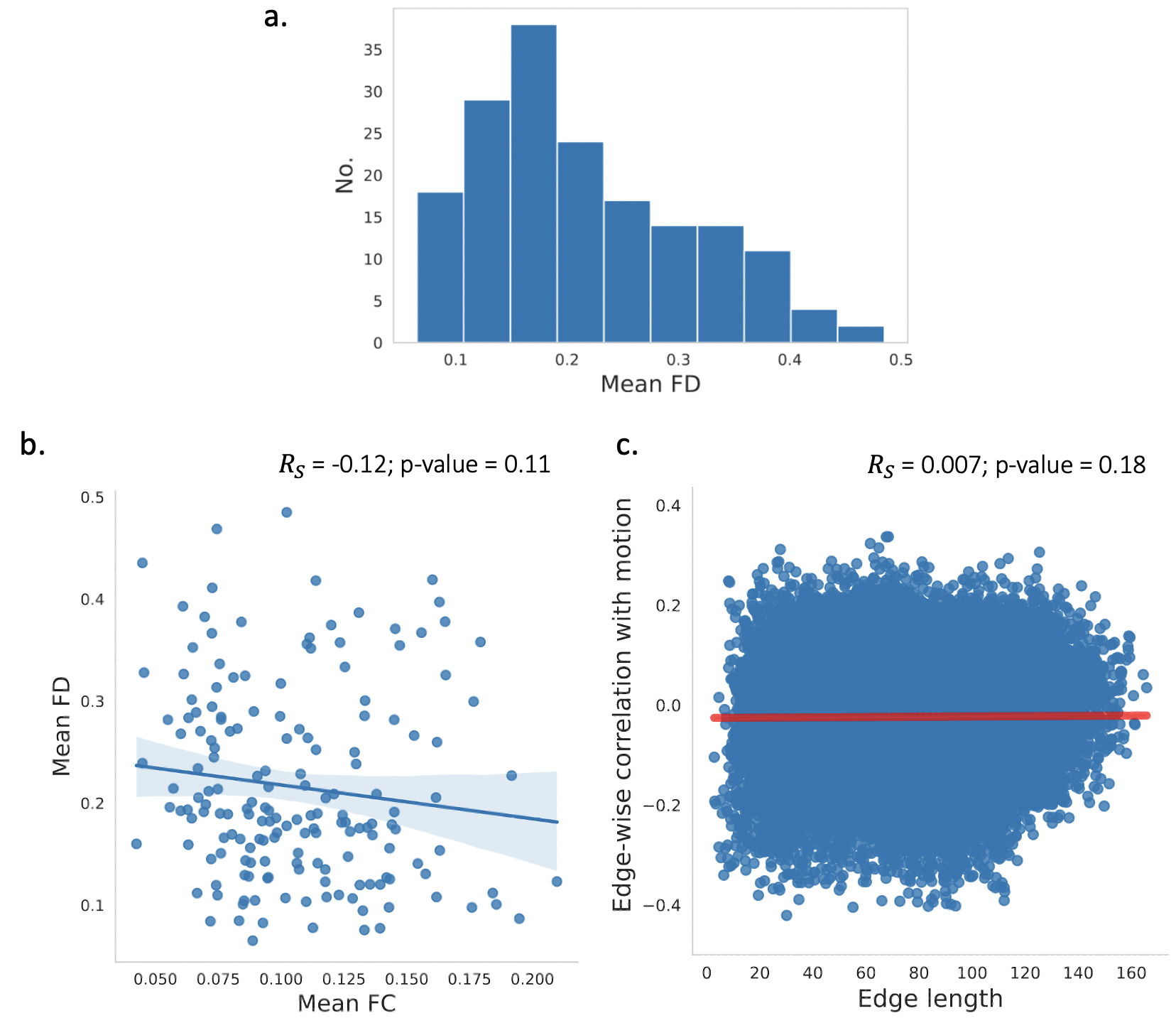

**FigS2**. a. Mean framewise displacement (Mean FD) distribution. b. Correlation between mean functional connectivity (Mean FC) and Mean FD. c. Distance dependence of motion. To calculate this, each edge was correlated with the Mean FD across subjects, and the resulting Spearman correlation coefficients were correlated with the edge length (i.e. Euclidean distance between a node pair forming an edge) for each edge.

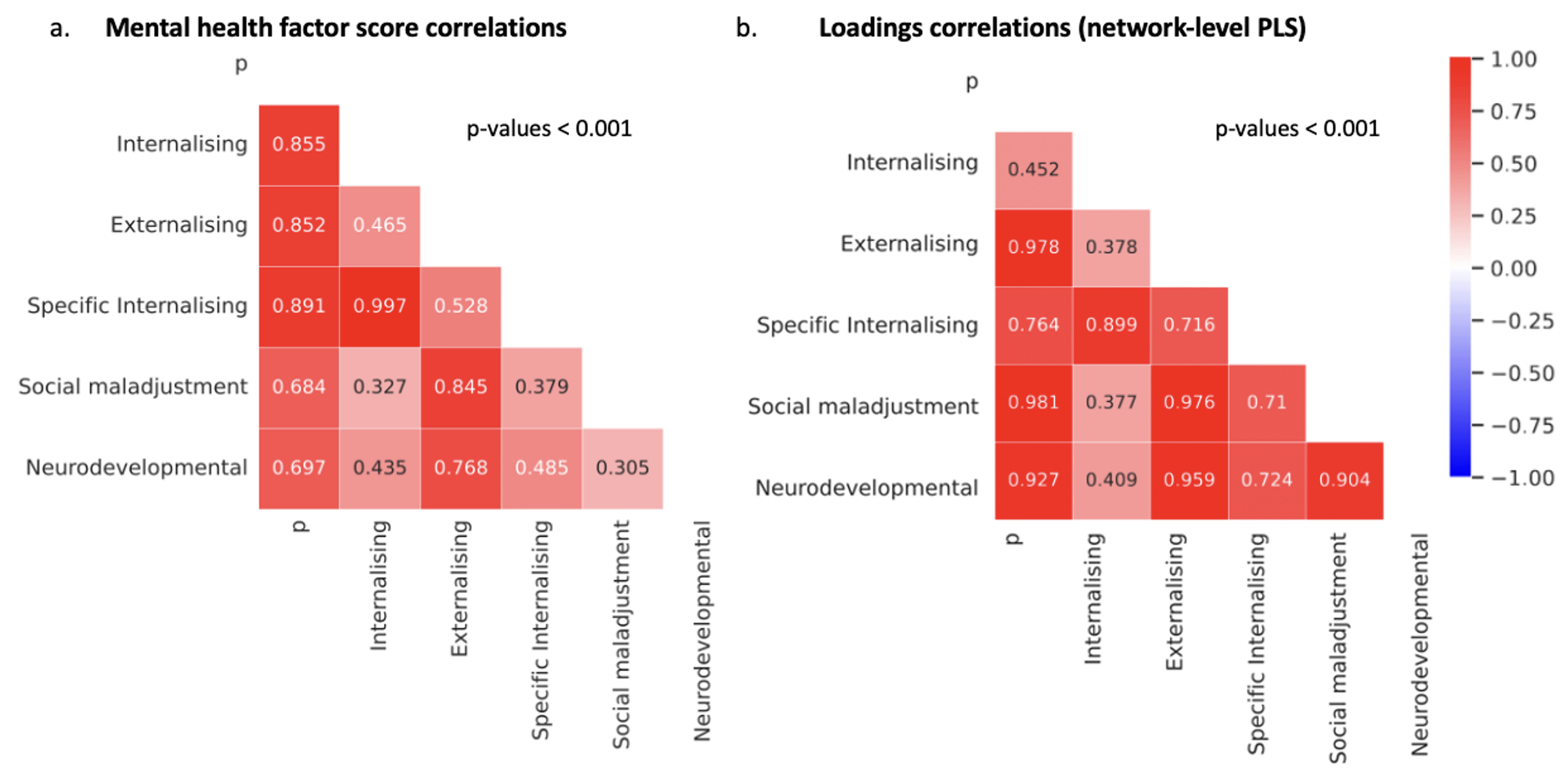

**FigS3**: a. Spearman correlation coefficients shown between each pair of mental health dimensions scores derived from the behavioural data, for the subset of participants with data on all mental health subscales and rs-fMRI (N = 67). b. Spearman correlation coefficients between the network-level connectivity loadings derived from PLSR analyses performed for each mental health dimension.

**Sensitivity analysis**

Several sensitivity analyses were performed to test the robustness of the results so far

with the smaller subset of participants with all mental health dimensions, as well as to

identify potential associations with age and sex.

The externalising, neurodevelopmental and social maladjustment dimensions derived from

the larger subset with externalising behavioural data (N = 771) were highly correlated

(R > 0.9) with their counterparts derived from the smaller participant subset with all

mental health dimensions (N = 378; FigS4). This indicated that the smaller subset

of participants (N = 378) offers a good level of statistical power for deriving the mental

health dimensions.

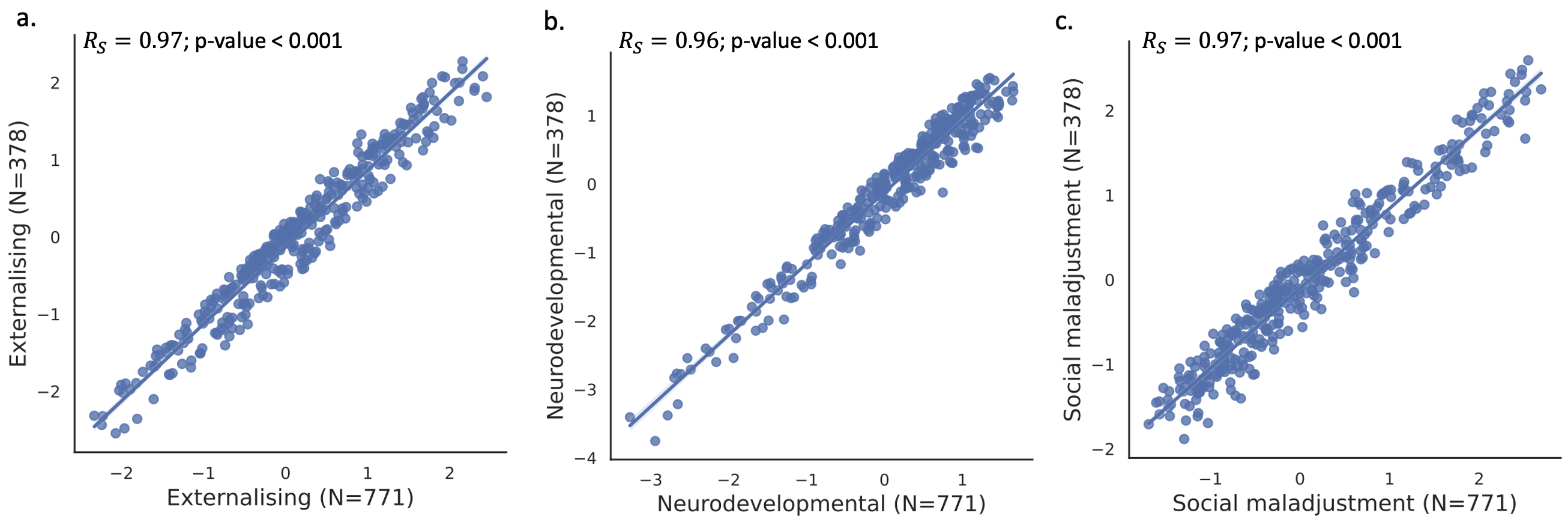

**FigS4**. Scatterplots of the Spearman correlations between the externalising mental health dimensions (externalising, neurodevelopmental and social maladjustment) derived from the subset of participants with data on the SDQ & Conners questionnaires (N = 771) and the externalising dimensions from the smaller subset of participants with data on the SDQ, Conners and RCADS questionnaires (N = 378), regardless of whether rs-fMRI data was available.

Next, the network-level PLSR analyses were repeated on the smaller subset of subjects

with all mental health dimensions and rs-fMRI data (N = 67). Largely similar patterns of network-level connectivity (FigS5; TableS1) to the initial results were identified. The variance explained by the network-level connectivity component was no longer significant, before FDR correction, for the externalising, neurodevelopmental and social maladjustment dimensions (TableS1), as opposed to the initial results with the larger subset (Table 2).

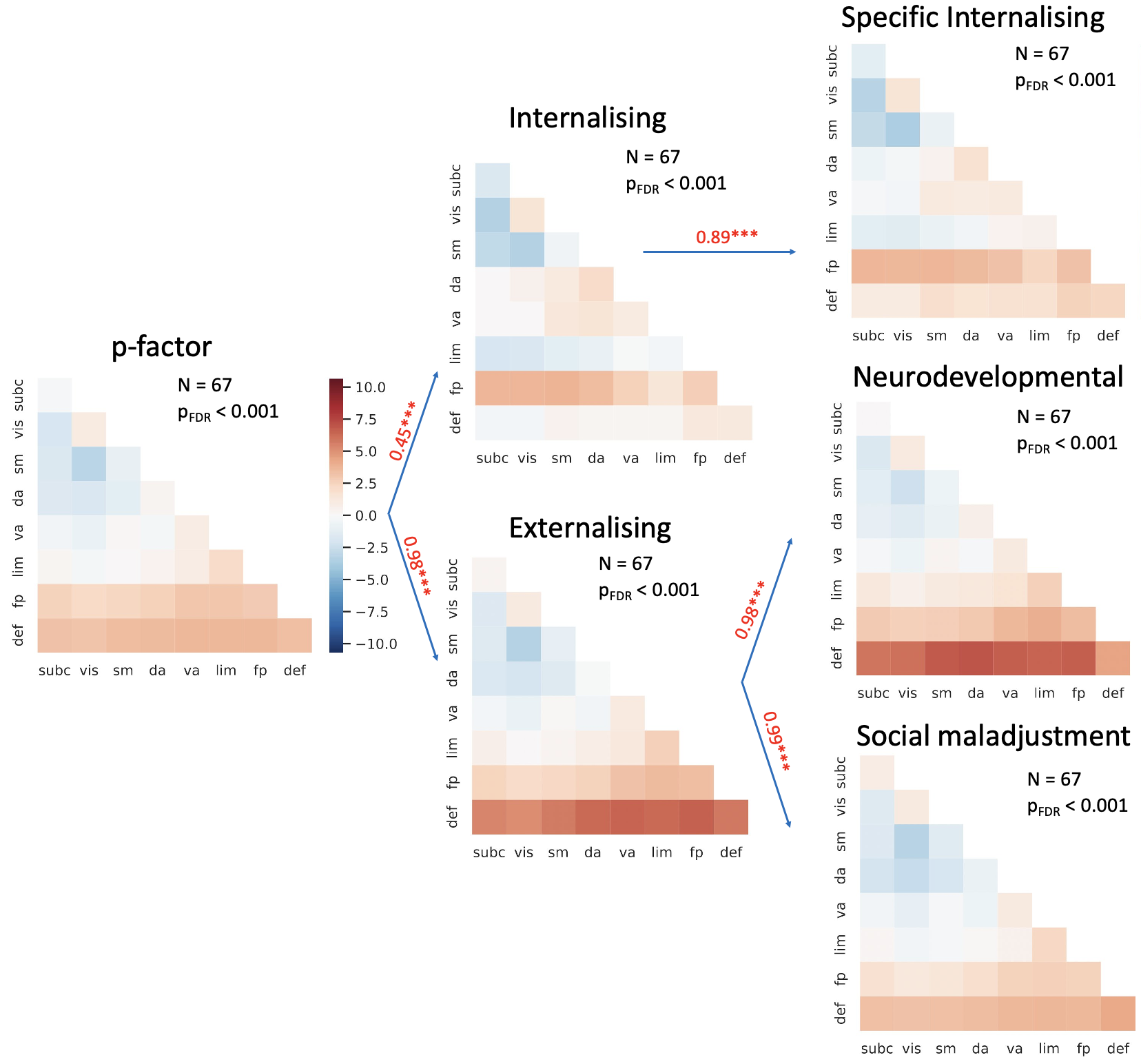

**FigS5**. Network-level functional connectivity x-loadings from PLSR analyses performed at the network level with each mental health dimensions in the ‘RCADS FC subset’ (N = 67). The loadings are shown for each of the 8 included ICNs (abbreviations: subc – subcortical, vis – visual, sm –somatomotor, da – dorsal attention, va – ventral attention, lim – limbic, fp – frontoparietal, def –default-mode network) on the diagonal, and between each pair-wise interaction between the 8 ICNs. The Spearman correlation coefficient between the loadings on the mental health dimensions is shown on each arrow. In red, the loadings which were positively correlated to each mental health dimensions are shown. The opposite trend is represented by the blue colours. The FDR-corrected *p-values* from the correlations between the x-scores and y-scores are also shown along with the number of subjects used for each of the six PLSR analyses.

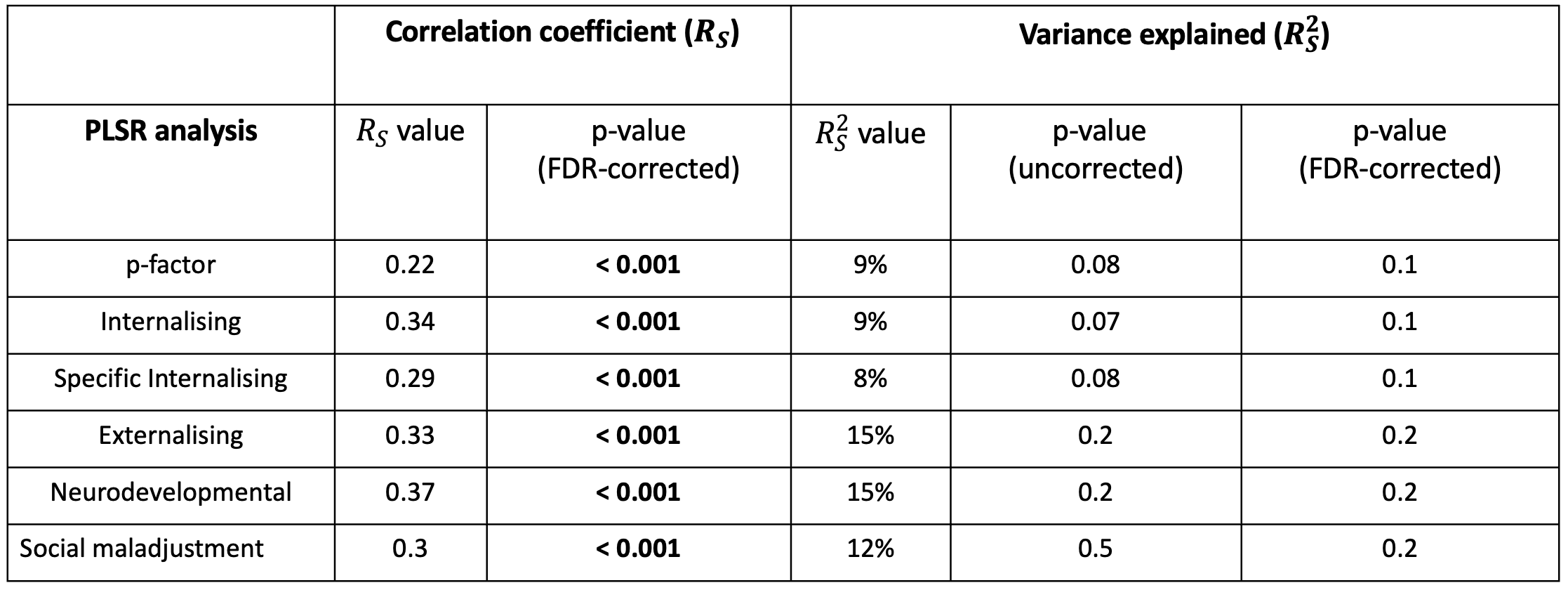

**TableS1**. Network-level PLSR results with each mental health dimensions in the ‘RCADS FC subset’ (N = 67). The Spearman correlation coefficient values between the PLSR-derived x-scores and y-scores for each of the six network-level PLSR analyses. The FDR-corrected p-values for these correlation coefficients are shown. The variance explained by the PLSR component from the corresponding network-level PLSR with each mental health dimensions. Significant p-values (*p-value* $\leq$ 0.05) are shown in bold.

The PLSR analyses were repeated without controlling for participant age and sex and the derived FC components (PLSR x-scores) were not significantly associated with these demographics, indicating robustness across age and sex differences.
